# MAPKAP Kinase-2 signaling enhances the resolution of colonic inflammation through specialized pro-resolving mediators

**DOI:** 10.64898/2026.10.13.758902

**Authors:** Hannah A. Bank, Lucia Suarez-Lopez, Xueyang Yu, Xavier de la Rosa, Tiffany Emmons, Yi Wen Kong, Paul W. Furlow, Eduardo J. Villablanca, Charles H. Cook, Charles N. Serhan, Michael B. Yaffe

## Abstract

Failure to resolve acute inflammation contributes to chronic inflammatory diseases, including inflammatory bowel disease, and inflammation-associated cancer. MAPKAP kinase-2 (MK2), a stress-activated effector of p38MAPK, is best known for promoting onset and amplification of inflammation through post-transcriptional regulation of cytokines. Here we identify an unexpected role for MK2 in coordinating resolution of intestinal inflammation. Following dextran sodium sulfate–induced colonic injury, MK2-deficient mice developed more severe, persistent inflammation, with sustained proinflammatory cytokine and chemokine production, prolonged neutrophil accumulation, and impaired macrophage recruitment during recovery. Myeloid-restricted MK2 deficiency recapitulated the impaired resolution phenotype. Lipid mediator metabololipidomics revealed MK2-deficient colons failed to undergo the transition from pro-inflammatory to pro-resolving lipid mediator production, exhibiting markedly reduced levels of specialized pro-resolving mediators (SPMs), including resolvins and lipoxins. Exogenous SPM administration during the recovery phase restored inflammation resolution in MK2-deficient mice, improving weight, disease activity, and mucosal healing. These findings establish a previously unrecognized role for MK2 in coordinating the transition from inflammatory injury to resolution and repair. Together with our prior finding that myeloid MK2 deficiency reduces colitis-associated tumor progression, these results suggest that inflammation-associated tumorigenesis depends not simply on inflammatory injury, but on its coupling to the resolution and tissue repair that follows.

## Introduction

Acute inflammation is an adaptive response to tissue injury or infection that attempts to restore homeostasis, largely mediated through the actions of neutrophils and macrophages (1–3). The inability to properly resolve an acute inflammatory insult plays a major role in the development of a variety of chronic inflammatory diseases and contributes to the development of multiple types of cancer (4–9).

Under normal conditions, tissue-resident macrophages and mast cells respond to tissue damage or infection by producing an assortment of cytokines and chemokines, resulting in an initial wave of neutrophil infiltration into the injured or infected tissue, followed by monocyte recruitment and differentiation into macrophages (10). Following successful elimination of pathogens and/or clearance of tissue debris, regeneration and repair ensues, restoring the tissue to some semblance of its pre-injured state, depending on the degree of scarring and fibrosis that occurs during the repair process. As tissues transition from a state of inflammation to a state of regeneration and resolution, there are concomitant changes in the patterns of gene expression seen in the infiltrating neutrophils and macrophages, which change from an initial pro-inflammatory neutrophil and M1-like macrophage state, respectively, to more reparative pro-resolving neutrophil and M2-like macrophage phenotypes (11–14).

Resolution of inflammation and the shift towards repair, historically thought to be a passive process (10,15), is now recognized as a highly dynamic active process largely mediated by several families of macrophage-derived lipids (3,10,16). These specialized pro-resolving lipid mediators (SPMs) include lipoxins, resolvins and protectins, all of which are produced by metabolic biosynthesis primarily from the eicosanoids arachidonic acid, eicosatetraenoic acid and docosahexaenoic acid (17,18).

Consistent with a model of active inflammation resolution, defective synthesis of pro-resolving lipid mediators has been postulated to underlie the pathogenesis of various inflammatory diseases, particularly in mucosal organs that are continuously exposed to a local microbiome (3,19,20). Patients with ulcerative colitis, for example, are reported to have low to absent levels of Lipoxin A_4_ (21), while reduced levels of lipoxins and/or Protectin D1 in the lungs are seen in humans with asthma and cystic fibrosis (22–24). Exogenous administration of SPMs or their precursors is therefore being actively explored as a strategy to suppress chronic inflammation and promote its resolution in a wide variety of disease contexts, including renal, pulmonary, and cardiovascular diseases (25,26). In preclinical studies for example, supplemental administration of either Lipoxin A_1_ or Resolvin E1 was shown to limit the extent of inflammation in murine models of inflammatory bowel disease from hapten-or DSS-induced colitis (27–29), while administration of Lipoxin A_4_ or B_4_ was shown to limit airway hyperresponsiveness and inflammation in murine models of asthma (23,30). Despite compelling pre-clinical efficacy, clinical translation of SPM therapy for inflammatory disease treatment remains at a relatively early stage (25,26) because of capital starvation.

There is growing interest in harnessing the inflammation-resolving activities of specialized pro-resolving mediators (SPMs) to prevent or limit cancer development and improve therapeutic responses (31–34). A relationship between inflammation and cancer can be traced back to the Edwin Smith Papyrus circa 3000 BC (35), although it was Virchow who first proposed that malignancies could arise at sites of chronic inflammation based on his observation of leukocyte infiltration into tumors (36). This concept was later revised by Dvorak’s influential description of tumors as “wounds that do not heal” (37), and subsequently expanded on by others (38). However, the mechanistic relationship between chronic inflammation, failed or dysregulated inflammation resolution, and cancer is complex and remains incompletely understood. Current evidence suggests that inflammation promotes tumorigenesis through a combination of genetic and epigenetic damage together with remodeling of the local tissue microenvironment into a niche that supports the survival, proliferation, immune evasion, and progression of premalignant and malignant cells (39–43). Dysregulation of inflammation resolution appears to contribute to this process by altering the recruitment, persistence, and functional programming of neutrophils and macrophages within the tissue microenvironment. This includes both prolonged activity of inflammatory, tissue-damaging myeloid cells and the inappropriate persistence or tumor-mediated co-opting of myeloid subsets with immunoregulatory and tissue-repair programs that promote angiogenesis, matrix remodeling, and tumor invasion (41,44–46). Whether defective production of specific SPMs is sufficient to act as driver of cancer development and progression, however, is less clear, particularly in human studies, though lower levels of lipoxinA_4_ and resolvin D1 in serum from patients with colon cancer has been reported (47,48).

Here we report an unexpected connection between the resolution of colonic inflammation, SPM production, and tumor development in mice lacking the stress-activated protein kinase MAPKAP Kinase-2 (MK2). MK2, a critical downstream effector kinase of the p38MAPK pathway, is best understood as a critical regulator of early inflammation onset and amplification by phosphorylating RNA-binding proteins that control the stability and translation of inflammatory mRNAs. This occurs through MK2-dependent phosphorylation of RNA-binding proteins that control the stability and translation of mRNAs encoding inflammatory cytokines such as TNF-α and IL-6 (49–52).

Using the dextran sodium sulfate (DSS) murine model of inflammation- and microbiome-mediated colitis-associated colon cancer (53), we and others previously reported that mice lacking MK2 showed a reduction in the growth of colon tumors following DSS treatment (54–56). The impairment in colon tumor growth in MK2 whole-body knockout animals was phenocopied in mice with myeloid-specific depletion of MK2, and shown to be a consequence of the failure of MK2-deficient macrophages to fully polarize into an M2-like state. This resulted in impaired macrophage production of several angiogenic growth factors (CXCL12, TIMP-1 and SERPIN E1/PAI-1), leading to impaired angiogenesis which slowed tumor development and progression (57).

We demonstrate that, despite reduced colon tumor progression, MK2-deficient animals are defective in inflammation resolution of their colitis as a consequence of delayed macrophage recruitment and defective SPM production in the inflamed colon. These findings reveal a previously unappreciated role for MK2 in regulating the resolution of inflammation, with implications for how the balance between persistent inflammation and tissue repair contributes to cancer development and progression in this model of colitis-associated cancer.

## Results

### MK2 Deficiency Exacerbates DSS-induced Acute Colitis

During the course of our previous work characterizing the effect of MK2 loss on DSS-induced colon cancer progression (56,57), we made the unexpected observation that the extent of intestinal inflammation induced by each cycle of DSS treatment – the stimulus used to induce colitis – appeared to be more severe in the MK2 knock-out animals, despite the fact that the MK2-deficient animals ultimately developed a reduced tumor cell mass.

In this study we further characterized the role of MK2 in regulating the onset and resolution of colon inflammation using the DSS model. Importantly, in this colitis model intraluminal DSS is used to disrupt the colonic epithelial barrier, resulting in submucosal microbial invasion and a potent mucosal inflammatory response (58). The model is particularly useful when studying the contribution of the innate immune system to the development of intestinal inflammation, since it is largely independent of T and B cells (59,60).

Phenotypic hallmarks of DSS-induced colitis include colonic shortening, weight loss, diarrhea, and rectal bleeding (53) which can be used to quantify the degree of colitis. Notably, we found that during the recovery period following five days of exposure to DSS-supplemented drinking water, MK2 whole body knock-out mice exhibited shorter colon length, sustained reductions in body weight, and higher rates of hematochezia, as indicated by the DAI score (Figure 1A-1D). The acute colitis in MK2-KO and MK2-WT mice was further characterized by quantifying the degree of mucosal injury (ulcers or crypt denudation), frequency of immune follicles (immune infiltrates), and dysplastic crypts (hyperproliferative glands surrounding ulcers). All three of these indicators of inflammation were more prevalent in MK-KO mice exposed to DSS (Figures 1E-F). Overall, when compared to MK2-WT mice, MK2-KO mice exhibited more severe DSS-induced colon inflammation marked by greater persistence of inflammation during the recovery period, suggesting that MK2 might play an important role during inflammation resolution.

**Figure 1.**
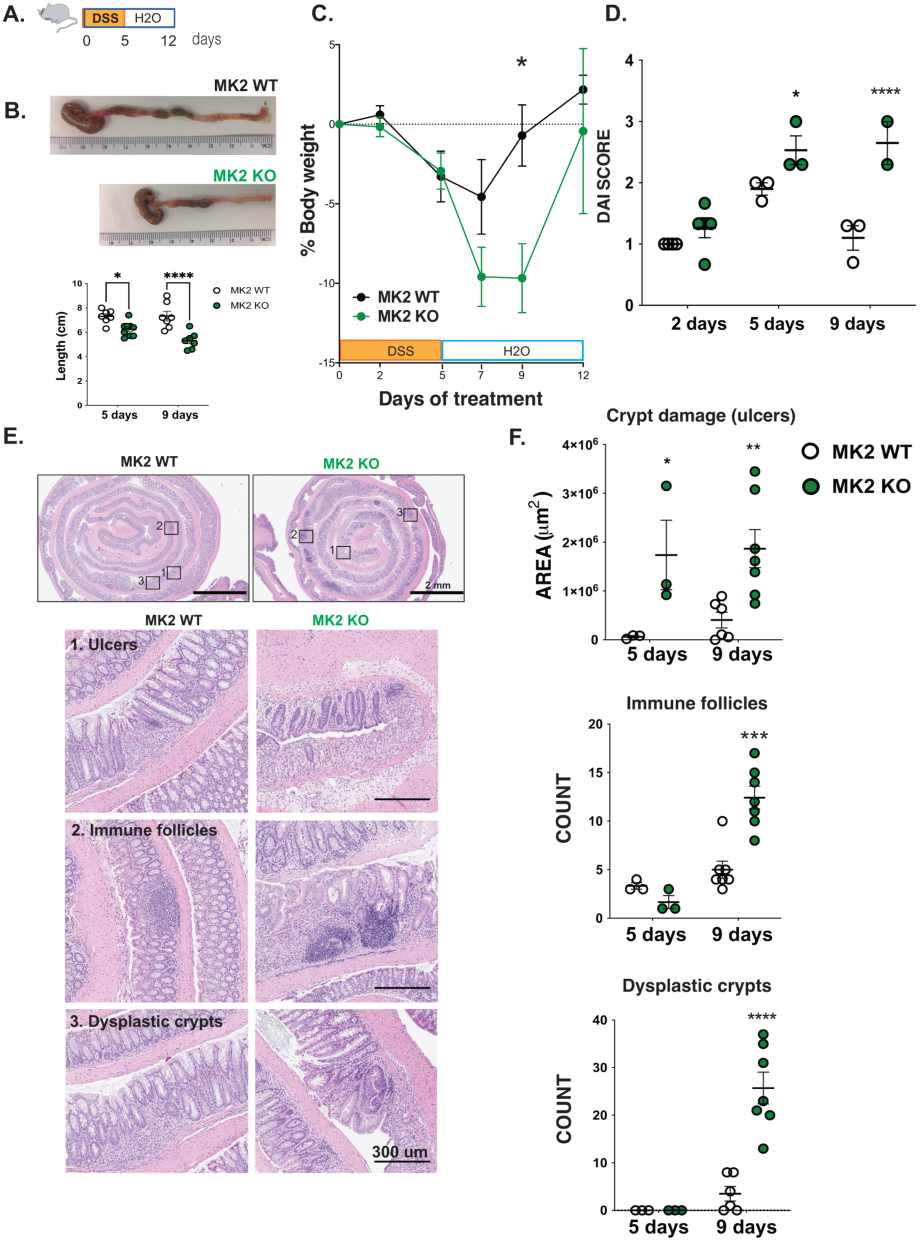
MK2 deficiency potentiates DSS colitis. **(A)** 2.5% DSS was administered in the drinking water for 5 days and mice allowed to recover for an additional 7 days. **(B)** Mice were euthanized at days 2, 5 and 9 of treatment and colon length measured. Representative pictures of colons at day 9 are shown. **(C)** Body-weight, recorded every 2-3 days during and following DSS treatment in MK2 WT and KO animals. **(D)** Disease activity index (DAI) was calculated by scoring weight loss, stool consistency and blood presence in feces. **(E)** Colon swiss rolls were prepared for pathology examination. Representative images from day 9 time point are shown. **(F)** Number of immune follicles, dysplastic crypts and ulcer extension were blindly measured in ImageScope. All plots in panels B, C, D, and F show data from 7-10 mice per group with mean ± SEM indicated. *p-value < 0.05; **p-value < 0.01; ***p-value < 0.001, ****p-value<0.0001; Panels C, D: 2-way ANOVA, Bonferroni post-hoc test. Panels B, F: Student’s t-test WT vs KO).

### MK2 differentially modulates the levels of pro- and anti-inflammatory cytokines and influences neutrophil and macrophage recruitment during the inflammation and resolution phases of acute colitis

To further assess the effect of MK2 loss on colon inflammation and resolution over time, we characterized the tissue cytokine profiles of MK2-WT and MK2-KO colons after DSS administration. Tissue homogenates prepared from the colons of WT and MK2 knock-out animals prior to treatment (day 0), after 5 days of DSS administration, and at day 9 in the middle of the recovery period, were analyzed for the expression of cytokines and chemokines involved in inflammation and tissue repair (Figure 2 and Supplemental Tables 1 and 2). Markedly, time-dependent, differences were observed in tissue cytokine/chemokine levels between the wild-type and MK2 whole body knock-out animals, both at the time of maximal tissue injury, and during the period of inflammation resolution. Compared to MK2-WT mice, MK2-KO mice displayed a sustained increase in levels of pro-inflammatory cytokines such as IL-6, IL-17, and LIF, and a decrease in the levels of anti-inflammatory repair-oriented cytokines including IL-10, IL-13, and M-CSF during recovery from DSS treatment (Figure 2A, B), indicative of a prolonged early phase of acute inflammation. Notably, all three cytokines have been implicated in promoting regulatory or repair-associated macrophage states and in restraining classical pro-inflammatory responses (11,61–64).

**Figure 2.**
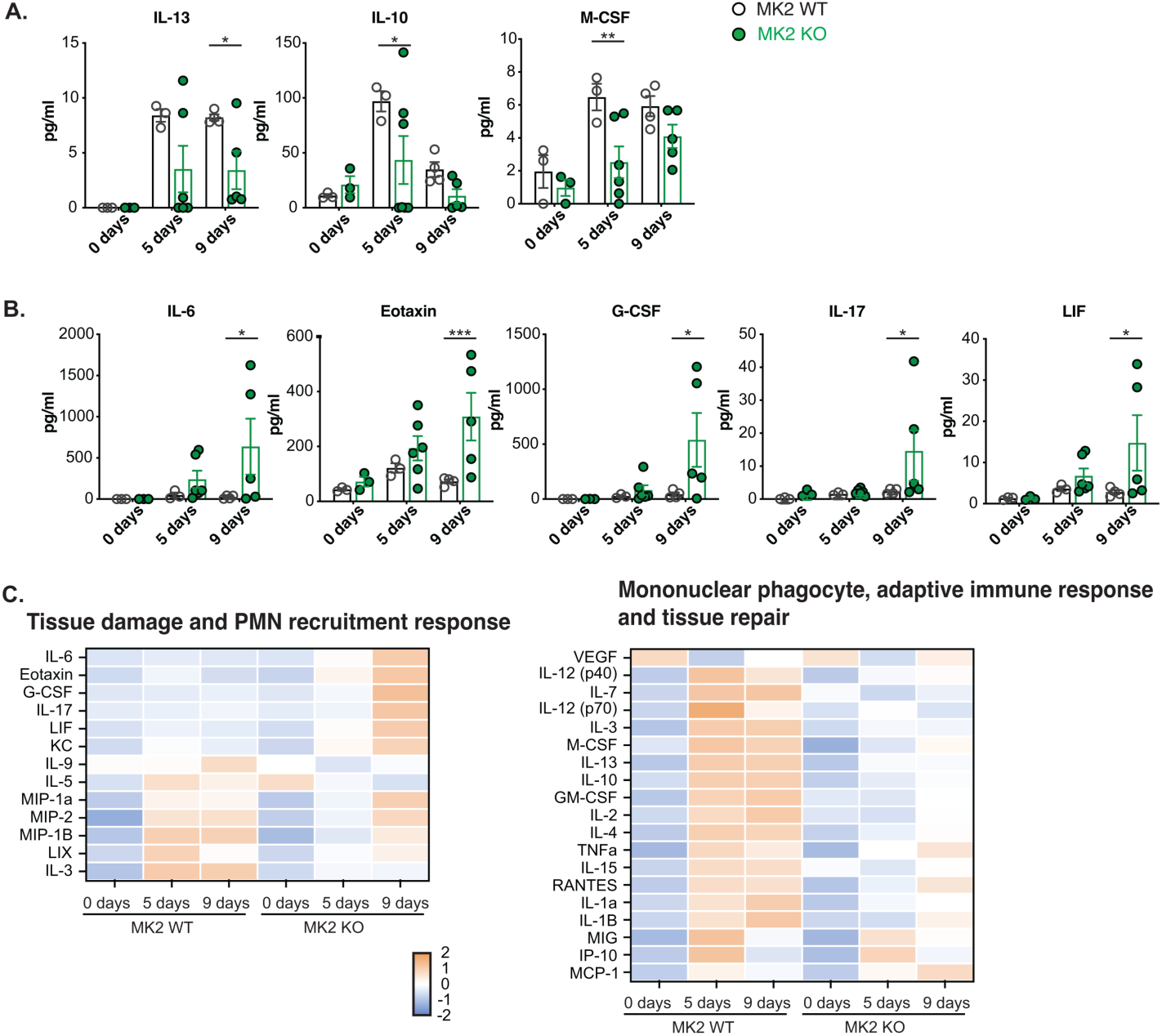
Altered cytokine levels associated with neutrophil and macrophage recruitment, tissue injury and repair in MK2-deficient versus MK2-proficient colons following DSS colitis. **(A, B)** Levels of representative anti-inflammatory/tissue repair (**A**), or pro-inflammatory (**B**) cytokines from DSS colons in MK2 KO mice compared to WT controls. **(C)** Heat map showing 32-plex Luminex quantification of murine cytokines/chemokines in colonic tissues of MK2 KO mice and littermate controls at days 0, 5 and 9 of DSS challenge experiment. Normalized z-scored values by analyte are shown here using the absolute numerical values shown in Supplementary Table 1. *p-value<0.05; **p-value<0.01; ***p-value < 0.001 using Student’s t-test WT vs KO.

Profiling of DSS-treated colons further revealed that cytokine/chemokine family members known to be involved in neutrophil recruitment were upregulated in MK2-KO mice exposed to DSS, whereas cytokine/chemokine family members involved in monocyte/macrophage recruitment and differentiation were down-regulated (Fig. 2C). For example, G-CSF and KC/CXCL1, which directly promote neutrophil production and recruitment, respectively, together with IL-17 and IL-6, which can amplify neutrophilic inflammation through induction of granulopoietic factors, chemokines, and endothelial activation, were upregulated 9 days following the onset of DSS administration, as was eotaxin, a chemotactic factor for eosinophils with some activity towards basophils and mast cells. Conversely, M-CSF, a major regulator of monocyte/macrophage differentiation, proliferation, and survival, showed reduced levels in the MK2-KO colons, as were the myeloid-associated cytokines IL-1β, IL-10, and IL-12p70/p40 (Figure 2C). Collectively, this cytokine profile was consistent with altered accumulation and/or activation of mononuclear phagocytes during recovery. Since that transition between early and late phases of acute inflammation is marked by infiltration of mononuclear phagocytes into a neutrophil-rich environment for the purpose of resolving inflammation, the observed pattern of dysregulated cytokine expression in MK2-KO mice suggested that MK2’s role in mediating the resolution phase of acute inflammation could be related to promoting macrophage recruitment.

To further investigate this, we analyzed the immune cell profiles of DSS-treated MK2-WT and MK2-KO colons over time by flow cytometry using the gating strategy shown in Figure 3A. Neutrophils and inflammatory monocytes within the CD45+ population were identified based on expression of CD11b, Gr1 and Lys6C, while activated tissue macrophages were identified in the CD45+ population by expression of CD64 (FcγR1), an activating receptor that signals through the ITAM-bearing FcR γ-chain in the setting of immune activation to facilitate immune effector functions (65). Notably, the MK2-KO animals displayed both an increase in the number of neutrophils in the colonic tissues at the end of the acute inflammation phase following 5 days of DSS treatment (Figure 3B), and a decrease in the number of activated tissue macrophages during the subsequent phase of inflammation resolution 4 days later (9 day time point) (Figure 3C).

**Figure 3.**
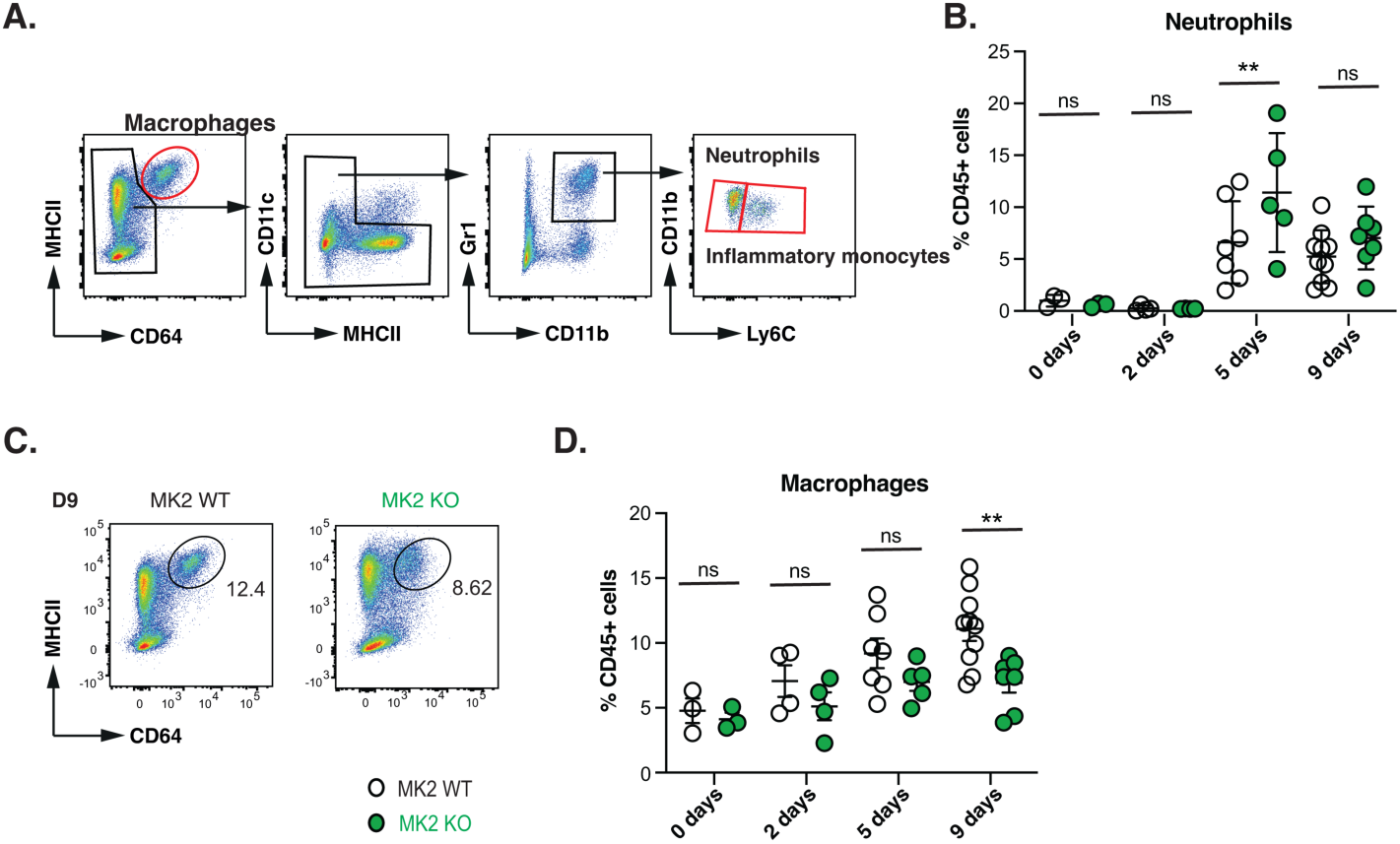
MK2 KO colon tissues have increased neutrophils following acute DSS colitis and reduced levels of tissue macrophages during resolution. Flow cytometry of colon tissues from MK2-KO and WT mice was performed at naïve state (day 0) and at days 2, 5, and 9 following DSS administration. **(A)** Gating strategy for macrophages, neutrophils, and Ly6C^hi^ monocytes from colon tissue, pre-gated on CD45+ singlets. **(B)** Frequency of neutrophils and Ly6C^hi^ monocytes at days 0, 2, 5, and 9, with mean ± SEM indicated. **(C)** Representative flow cytometry plots of colon MK2 WT or KO macrophages at day 9. (**D**) Frequency of colon macrophages at days 0, 2, 5, and 9 with mean ± SEM indicated. *p-value<0.05; ***p-value<0.001 using Student’s t-test WT vs KO.

### Myeloid MK2 is responsible for mediating resolution of inflammation in acute colitis

We next investigated whether the persistent late-stage inflammation and failure of colitis resolution that we observed in whole body MK2 knock-outs (Figure 3) was specifically due to could be similarly attributed to defective myeloid MK2 signaling. To examine this we used LysM-Cre mice to knock out MK2 in the myeloid compartment (MK2-LysM-KO) and treated the MK2 myeloid-specific knockout animals using the same DSS regimen described above. Colons from MK2-LysM-KO mice exposed to oral DSS recapitulated the phenotypic changes seen in colons from the global MK2-KO mice. Specifically, MK2-LysM-KO mice displayed shorter colon length, sustained reductions in body weight, increased hematochezia, and higher prevalence of mucosal injury, immune follicles, and dysplastic crypts when compared to animals with intact MK2 (Figure 4A, B). Cytokine profiling of colonic tissue homogenates from DSS-treated MK2-LysM-KO animals showed persistent expression of proinflammatory mediators (IL-6, Eotaxin, G-CSF, IL-17, and KC) compared to the MK2-LysM-WT animals, particularly at day 9 in the middle of the recovery period (Fig 4C, Supplemental Tables 1 and 2). These changes closely mimic what was observed in the MK2 whole-body knock-outs (Figure 2B, Supplemental Tables 1 and 2). Furthermore, immune cell profiling of the colons from DSS-treated myeloid-specific MK2 knockouts showed a similar reduction in tissue macrophages, when compared to MK2 wild-type controls, at days 5 and 9 days (Figure 4D), again recapitulating the results seen with the whole body MK2 knock-outs. There were no significant differences in total monocyte or neutrophil numbers between MK2-LysM-KO and MK-LysM-WT colons (Figure 4D). Taken together, these data indicate that the failure to resolve acute colonic inflammation in MK2 deficient animals results from loss of MK2 activity in the myeloid compartment, leading to impaired recruitment of tissue macrophages.

**Figure 4.**
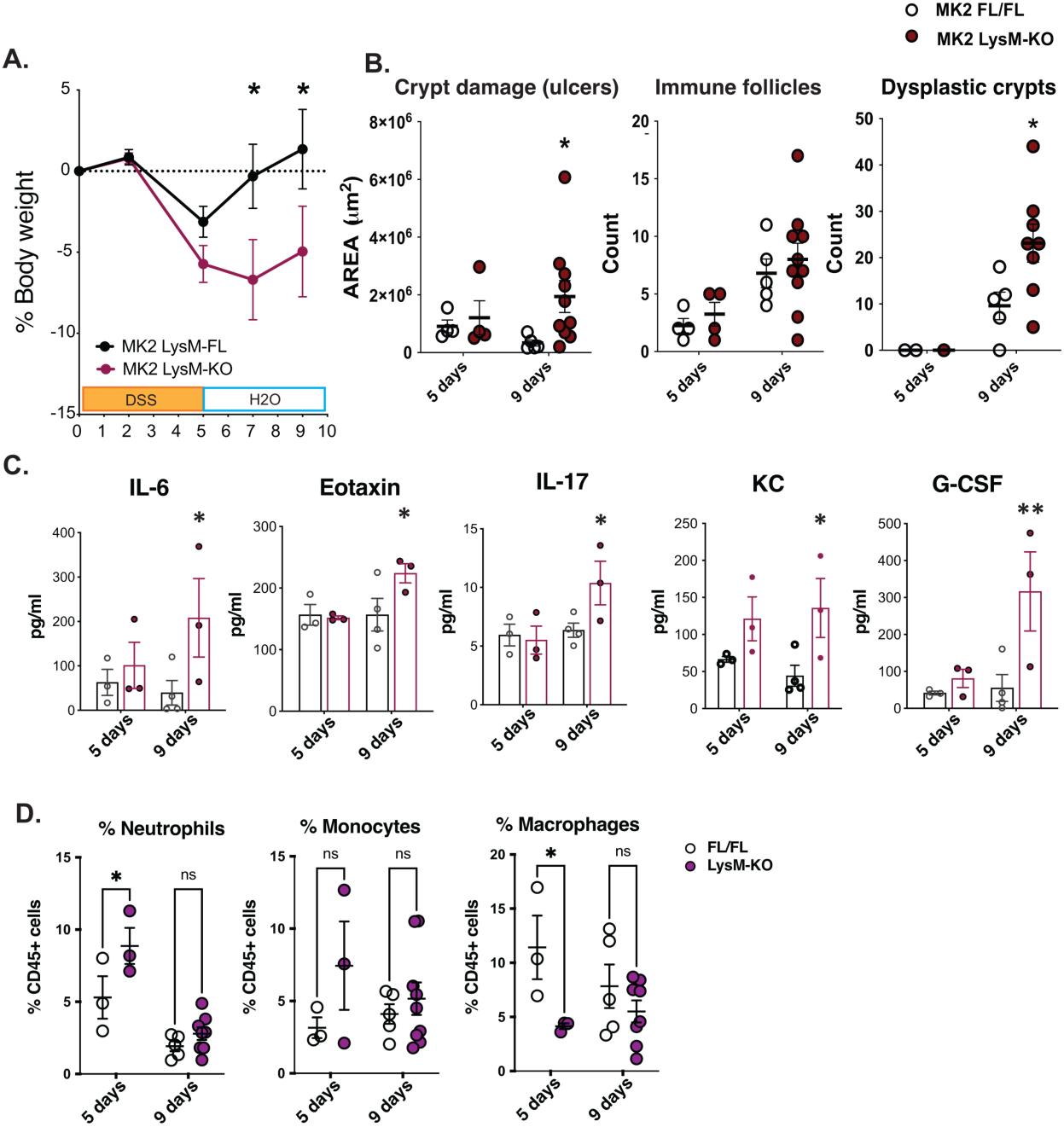
Myeloid-specific depletion of MK2 recapitulates colitis resolution failure. **(A)** 2.5% DSS was administered in the drinking water for 5 days and mice allowed to recover for an additional 7 days. Weight was recorded every 2-3 days. **(B)** Colon swiss rolls were prepared for pathology examination. Ulcer extension, number of immune follicles, and dysplastic crypts were blindly measured in ImageScope. **(C)** Levels of representative up-regulated cytokines in MK2 LysM-KO mice compared to littermate controls. **(D)** Frequency of macrophages, neutrophils and Ly6C^hi^ monocytes at days 5 and 9 post DSS challenge analyzed by flow cytometry. Plots in panels A-D indicate data from 3-10 mice per group with mean ± SEM indicated. *p-value < 0.05; **p-value < 0.01; ***p-value < 0.001; ****p-value<0.0001; ns indicates not significant. Panels A: 2-way ANOVA, Bonferroni post-hoc test. Panels B, C and D: Student’s t-test LysM-FL vs LysM-KO).

### MK2 promotes pro-resolving SPM production during acute colitis

To investigate the role of MK2 in local SPM production during acute colitis, we measured lipid mediator (LM) profiles in the colons of MK2-WT and MK2-KO mice 5 and 12 days after DSS exposure. Using targeted LC-MS/MS-based LM metabololipidomics, a wide range of SPMs were assayed (full list with stereochemistries in Supplemental Table 3), including D-series resolvins (RvDs), protectins (PDs), and maresins (MaRs) from the DHA metabolome as well as E-series resolvins (RvEs) from the EPA metabolome, and lipoxins (LX) from the AA metabolome (Figure 5, Supplemental Tables 4 and 5). Pro-inflammatory mediators, such as prostaglandins (PGs), thromboxane (TX), and LTs, were also measured from the AA metabolome (Supplemental Tables 4 and 5).

**Figure 5.**
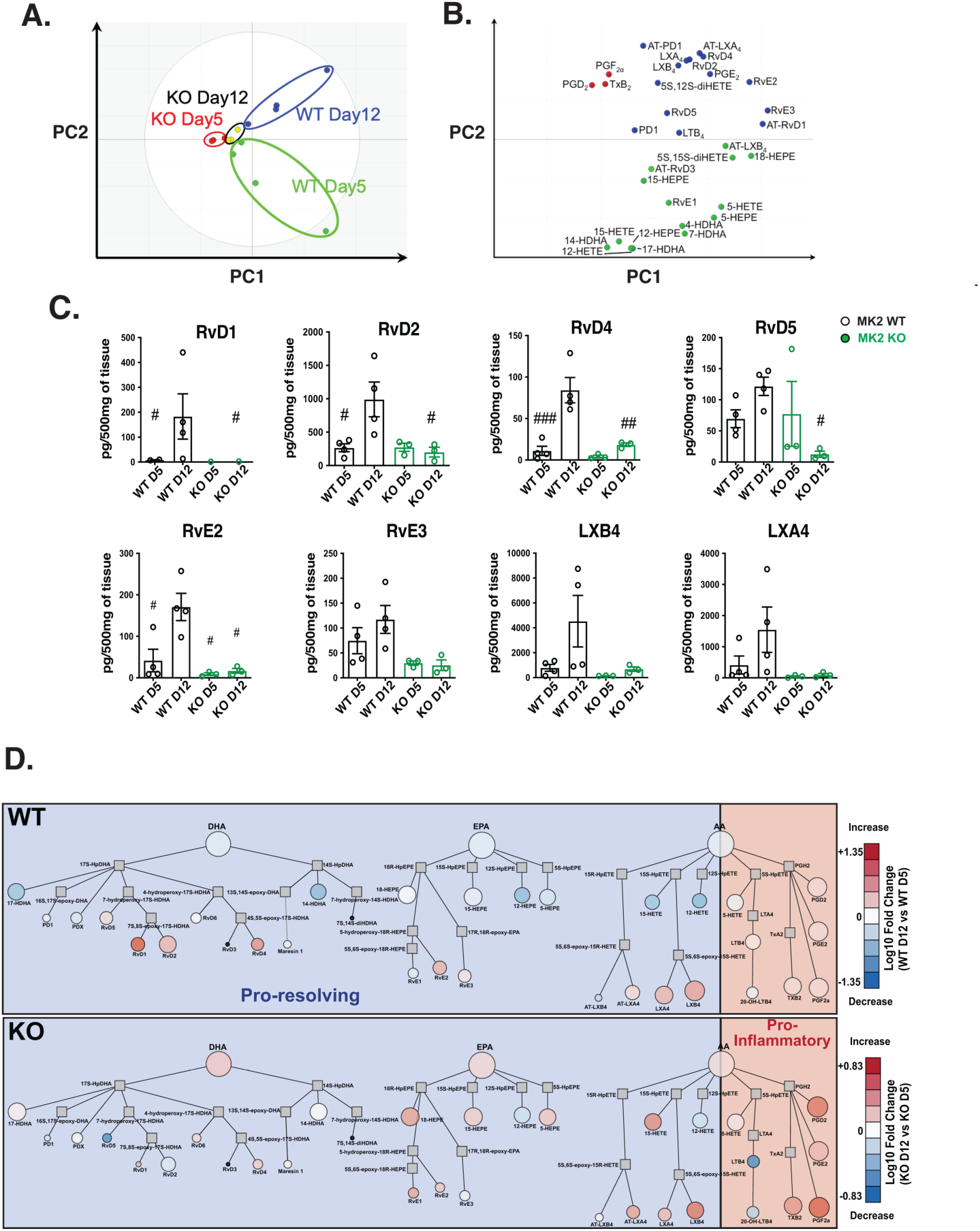
MK2 promotes inflammation-resolving SPM production during acute colitis. Colons from MK2-WT and MK2-KO mice were harvested on days 5 and 12 following DSS-colitis and analyzed for lipid mediators using LC-MS/MS metabololipidomics. **(A-B)** Principal component analysis showing temporal clustering of mediators from WT and KO animals at day 5 and day 12, in LM species-associated, two-dimensional score plot **(A)** and two-dimensional loading plot **(B). (C)** Quantitation of RvD1, RvD2, RvD4, RvD5, RvE2, RVE3, LXB4, and LXA4 in colons after DSS-induced colitis at days 5 and 12. Plots show data from n=3-4 individual animals per group with mean ± SEM indicated. #p<0.05; ##p<0.01; ###p<0.001 vs. WT day 12. n.s. indicates not significant, using one-way ANOVA followed by Tukey’s multiple comparisons test. **D)** LM Network visualization of WT vs. KO mice colons after DSS-colitis at day 12 vs. day 5. Node size indicates absolute amount of each metabolite in pg on day 12, using a log_10_ scale. Node color indicates log_10_-fold change in amount during resolution (day 12 vs day 5). Node shape: circles indicate measured metabolites, squares indicate non-measured pathway intermediates. Small black circles indicate not detected. Pro-resolving metabolites are shown in the blue shaded box, pro-inflammatory metabolites are shown in the red shaded box. Diagram created using Cytoscape **3.10.4** software.

Principal component analysis (PCA) using colonic LM levels from mice with DSS-colitis demonstrated separate temporal clustering of the samples at days 5 and 12 in the MK2-WT animals, which largely separated along both PC1 and PC2 compared to the MK2-KO samples (Figure 5A). In contrast, samples from the colons of MK2-KO animals at days 5 and 12 showed very little spatial separation along either PC axis on the scores plot. Examination of the PC loadings plot shows that pro-inflammatory lipid mediators (i.e. PGD_2_, TxB_2_) largely separate from pro-resolving lipid mediators (i.e. RvE2, RvE3, AT-RvD1, 18-HEPE) along PC1, whereas the upstream, precursors of pro-resolving lipid mediators (i.e. 17-HDHA) largely separate from the downstream mature pro-resolving lipid metabolites (i.e. RvE2, RvE3, LXA_4_) along PC2. The temporal trajectory of the WT samples on the scores plot in PC space correlated with a transition from pro-inflammatory to immature pro-resolving lipids, and then to mature pro-resolving lipids on the loadings plot, whereas no such transition was observed in the MK2 KO samples. Notably, there was a 3-fold or greater increase in the mean values of SPMs associated with inflammation resolution, (RvD1, RvD2, RvE2, and LXA_4_) from days 5 to 12 in the colons of DSS-treated MK2-WT mice that was not observed in the colons of MK2-KO animals (Figure 5C). Interestingly, LXA₄ and LXB₄, which were largely absent from the colons of MK2-KO animals, are endogenous counter-regulatory lipid mediators that limit neutrophil adhesion and transendothelial migration and attenuate leukotriene-driven inflammatory signaling (3,66,67), consistent with the increased presence of neutrophils at 5 days and pro-inflammatory cytokines at 9 days observed in the MK2 knockout animals. Taken together, these findings implicate MK2 in the mechanism by which acute colonic inflammation transitions from a pro-inflammatory phase to an anti-inflammatory/wound resolution phase, through an SPM-dependent process.

### SPM administration in acute colitis rescues the inflammation resolution defect in MK2-deficient mice

To directly test whether the lack of SPM production in MK2-deficient animals was responsible for the more severe colitis phenotype, we examined whether administration of supplemental SPMs could rescue the defect in inflammation resolution observed in MK2-KO mice during the recovery phase of DSS colitis. A mix of pro-resolving SPMs (LXA_4_, RvD1, RvD2, RvD4 and RvE2), or DMSO vehicle control, was administered by IP injection every other day beginning on day 5 following DSS administration. While the MK2 KO mice treated with either DMSO or the SPM cocktail showed similar body weight loss on day 7, the MK2-KO mice treated with SPMs regained their weight by day 9 significantly more efficiently than those receiving vehicle (Figure 6A).

**Figure 6.**
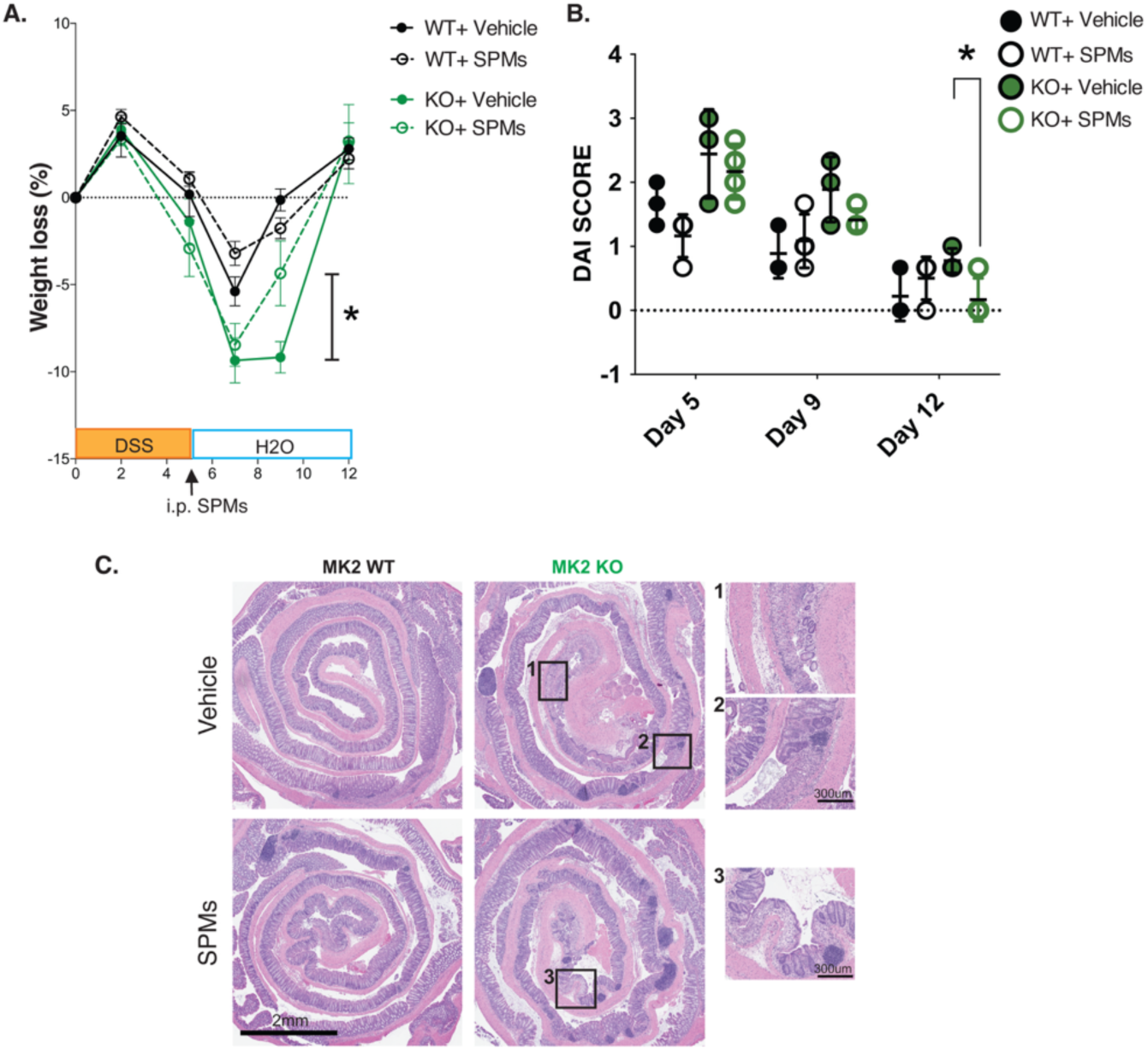
Supplemental SPM administration rescues inflammation resolution in MK2 KO mice with DSS-induced colitis. **(A)** 2.5% DSS was administered in the drinking water for 5 days prior to recovery for an additional 7 days. During the recovery phase, mice received every-other-day intraperitoneal SPMs (LXA4, RvD1, RvD2, RvD4 and RvE2) or vehicle control from days 5 to 12. Weight was recorded every 2-3 days. **(B)** Disease activity index (DAI) was calculated by scoring weight loss, stool consistency and blood presence in feces at Day 5, 9 and 12. **(C)** Colon swiss rolls were prepared for pathology examination. Representative images from day 9 time point are shown. Magnified squares show reduced ulcerated mucosal areas in SPMs-treated [3] compared to vehicle-treated MK2 KO mice in [1] and [2]. Plots in panels A and B indicate data from 2-4 mice per group with mean ± SD indicated. *p-value<0.05 using Student t-test, KO Vehicle vs KO + SPM.

Most strikingly, by day 12, the disease activity index in the MK2-KO mice treated with SPMs was significantly lower than in the MK2-KO mice treated with the vehicle control and was now comparable to that of MK2-WT littermates whose inflammation had fully resolved (Figure 6B). Colon tissue from SPM-treated MK2-KO mice also showed reduced mucosal ulceration at day 9 compared to vehicle-treated MK2-KO mice (Figure 6C). Overall, SPM administration was sufficient to restore inflammation resolution in MK2 knock-out animals following DSS-induced colitis.

## Discussion

Our findings identify an unexpected role for MK2 in coordinating the transition from acute intestinal inflammation to tissue resolution and repair. Although MK2 is conventionally viewed as a pro-inflammatory kinase because of its ability to stabilize and promote the translation of cytokine transcripts (49,50,68,69), MK2-deficient mice did not exhibit an attenuated response to DSS-induced epithelial injury. Instead, they developed persistent mucosal inflammation characterized by sustained expression of neutrophil-associated cytokines and chemokines, reduced expression of factors supporting macrophage differentiation and repair-oriented functions, and markedly diminished production of pro-resolving lipid mediators. These findings suggest that MK2 is required not only for the initiation and amplification of inflammatory signaling, but also for the subsequent reprogramming of the tissue environment that terminates neutrophil-dominated inflammation and permits macrophage-dependent clearance and mucosal repair.

This dual function may help reconcile apparently conflicting results regarding inhibition of p38–MK2 signaling in intestinal inflammation. Activation of p38 and MK2 is prominent in human inflammatory bowel disease and in experimental colitis, and pharmacological inhibition of this pathway can reduce production of TNFα and other inflammatory mediators, in the colons of human patients (70,71) and in a murine T-cell transfer model of IBD (72), culminating in fewer pathological findings. In contrast, whole-body MK2 deletion was found to exacerbate mucosal inflammation in a murine TNF-α overexpression model of IBD (73), indicating that the consequences of inhibiting this pathway may depend on the cellular compartment, phase of the inflammatory response, and duration of pathway inhibition. Consistent with this context dependence, fibroblast-restricted disruption of MK2 signaling was recently shown to reduce inflammation and intestinal fibrosis in a DSS-colitis model (74), whereas our results demonstrate that global or myeloid-restricted MK2 deficiency compromises the coordinated resolution of acute epithelial injury. Thus, MK2 cannot be classified simply as a unidirectional driver of intestinal inflammation. Rather, its effects appear to change over the course of the inflammatory response and among epithelial, immune, and mesenchymal compartments.

A particularly informative finding was that MK2-deficient mice experienced more severe and persistent inflammation yet developed less extensive tumor progression (56,57). This apparent paradox argues that the magnitude or duration of inflammation alone does not determine the tumor-promoting consequences of intestinal injury. In the AOM/DSS model, carcinogen-initiated epithelial cells are subjected to repeated cycles of DSS-induced injury followed by periods of recovery (75). Inflammatory mediators generated during the injury phase can cause additional genomic damage and provide survival signals to initiated cells, but the recovery phase supplies a complementary set of tumor-promoting conditions: epithelial proliferation, stem-and progenitor-cell expansion, angiogenesis, extracellular-matrix remodeling, and the accumulation of macrophages and fibroblasts that support tissue reconstruction. When appropriately terminated, these processes restore epithelial integrity. When repeated in tissue containing initiated cells, however, the same regenerative programs can facilitate the survival, selection, and expansion of premalignant clones (41,76).

Our results therefore support a model in which MK2 promotes the transition from a neutrophil-dominated injury response to a macrophage-and stroma-supported reparative state. Loss of MK2 appears to trap the colon in a destructive, poorly resolving inflammatory phase: neutrophilic inflammation persists, pro-resolving lipoxins and macrophage-supporting signals are diminished, and effective epithelial and stromal repair is compromised. Although this state produces greater acute tissue damage, it may provide less of the organized regenerative environment required for the progressive expansion of AOM-initiated epithelial clones. In this interpretation, colitis-associated tumorigenesis is driven not simply by uninterrupted inflammation, but by repeated coupling of inflammatory damage to compensatory resolution and repair. The report that mesenchymal MK2–HSP27 signaling promotes intestinal tumorigenesis is consistent with this model and suggests that MK2-dependent stromal repair programs may also be among the mechanisms connecting recurrent injury to tumor progression (77). This interpretation does not exclude additional, non-mutually exclusive explanations for the reduced tumor burden in MK2-deficient mice. MK2 loss could directly impair the proliferation or survival of initiated epithelial cells, alter angiogenesis or extracellular-matrix remodeling, or eliminate trophic signals normally supplied by macrophages and fibroblasts. Severe unresolved injury might also remove initiated epithelial cells rather than permit their recovery and expansion. Nevertheless, the dissociation between inflammatory severity and tumor progression observed here argues against a simple linear model in which more inflammation necessarily produces more cancer.

Our findings supporting a complex relationship between the extent of inflammation and subsequent tumor development are in strong agreement with a study from Greten et al who explored cell type-specific knock-outs of IKKβ in this same DSS colitis-associated cancer model (78). In that study, epithelial-specific loss of IKKβ resulted in reduced tumor incidence despite greater histologic evidence of tissue damage and colonic ulcerations, indicating that the tumor reduction was not the result of reduced inflammation. Myeloid-specific loss of IKKβ in this colitis model reduced both tumor number and tumor size, which the authors interpreted as evidence of myeloid-derived paracrine factors that were required for tumor promotion and growth.

The relevance of these findings to human ulcerative colitis and other forms of inflammatory bowel disease should be considered in light of the strengths and limitations of the DSS model. DSS causes direct chemical injury to the colonic epithelial barrier, followed by microbial translocation and a predominantly innate inflammatory response (53,58–60). It reproduces important features of ulcerative colitis, including epithelial erosion, crypt injury, neutrophil accumulation, ulceration, and cycles of mucosal damage and restitution (79). Importantly, spatially distinct neutrophil subpopulations associated with the colonic epithelium or localized to the lamina propria in human UC patients have recently been shown to be faithfully reproduced in DSS-treated mice, revealing location-specific differences in activation state and function (79). However, the DSS model does not reproduce the complex genetic, adaptive-immune, microbiome, and environmental origins of human IBD. MK2 deficiency in the DSS model should therefore not be interpreted as a direct model of MK2 dysfunction in patients with IBD. Rather, the model reveals a potentially relevant biological principle: the extent of p38–MK2 signaling may help determine whether barrier injury progresses toward successful resolution or remains locked in a neutrophil-rich, tissue-damaging state.

Finally, these findings also suggest caution in considering MK2 inhibition as a treatment for chronic intestinal inflammation or colitis-associated cancer. Inhibiting MK2 during established inflammatory amplification might reduce selected cytokines or tumor-supportive stromal programs, whereas sustained or systemic inhibition could interfere with pro-resolving mediator production, macrophage-dependent clearance, and mucosal recovery. The therapeutic consequences are therefore likely to depend on timing, duration, and cellular target. Defining the separable functions of MK2 during injury, resolution, and repair may make it possible to suppress its chronic tumor-promoting activities without disabling the mechanisms required to restore intestinal homeostasis.

## Material and Methods

### Sex as a biological variable

Our study exclusively examined male mice in agreement with the STAR protocol recommendations for the DSS-colitis model (80). Studies involving the DSS-colitis and AOM DSS-colorectal cancer models have shown a protective effect of estrogens and decreased effect size in female mice (80–82) and thus male mice are generally used for this model in the literature.

### Mice

MK2 KO mice were generated as described previously (Suarez-Lopez et al., 2018) and maintained in a C57BL/6N background. For the generation of myeloid specific MK2 knockout animals (LysM-KO mice), MK2 homozygous floxed mice (MK2 FL/FL) were crossed with the myeloid specific *Lyz2-Cre* line(83). LysM-KO and non Cre-carrier littermate controls (LysM-FLFL) were backcrossed 5 generations to C57BL/6N.

### Acute DSS colitis

8-12 weeks old male mice were administered 2.5% DSS (MP biochemicals) in the drinking water for 5 days to induce acute colitis. Disease activity index was measured every 2 days. The DAI is the combined score of weight loss compared to initial weight, stool consistency, and bleeding. Scores were defined as follows: weight loss: 0 (no loss), 1 (1-5%), 2 (5-10%), 3 (10-20%), and 4 (>20%); stool consistency: 0 (normal), 2 (loose stool), and 4 (diarrhea); and bleeding: 0 (no blood), 2 (Hemoccult positive), and 4 (gross bleeding, blood around anus).

All mouse studies described here were approved by the MIT Institutional Committee for Animal Care and conducted in compliance with the Animal Welfare Act regulations and other federal statutes relating to animals and experiments involving animals and adhere to the principles set forth in the Guide for the Care and Use of Laboratory Animals, National Research Council, 1996 (Institutional Animal Welfare Assurance No. A-3125– 01).

### Pathology evaluation of colon tissues

Intestinal tissues were prepared in swiss rolls for pathology evaluation. Briefly, rolls were fixed overnight in neutral buffered formalin and paraffin embedded. 4µm slides were cut and H&E stained. Slides were scanned in a Leica slide scanner before analysis and ulcerated areas, number of dysplastic crypts and immune follicles per swiss role were blindly measured on ImageScope software (Leica).

### Cytokine analysis

Colon tissues were snap frozen and disrupted in a tissue homogenizer (Genogrinder). Tissue homogenates were then lysed in RIPPA buffer containing proteases/phosphatases inhibitors cocktails (Roche). 60μg of total protein per sample was sent to Eve technologies for multiplex based cytokine/chemokine quantification.

### FACS analysis of immune populations

Colons were flushed with cold PBS and weight of tissue processed for FACS was recorded for later normalization. Tissue was prepared for FACS as described previously(Jijon et al., 2017). Briefly, tissue was minced with a razor blade and digested with Liberase TL (0.3mg/ml)/DNAse (10μg/ml) solution for 40 min at 37°C. Digested tissue was then filtered through 100uM mesh and washed in PBS solution containing 1% FBS, 2mM EDTA. Peritoneal resident macrophages were obtained by peritoneal lavage using 5 mL of ice-cold PBS supplemented with 2% FBS. Blood was collected by cardiac puncture and leukocytes were recovered after RBC lysis. Spleens were as well collected and dissociated thoroughly between frosted glass slides. Splenocytes were recovered after red blood cells lysis and processed for FACS analysis.

Single cell suspensions were incubated for 10 min at room temperature with Fc blocking (CD16/32) antibody (eBioscience) prior to staining with fluorochrome-conjugated antibodies against mixtures of the following antigens: CD45, CD19, CD4, CD90.2, CD127, CD3, CD25, NK1.1, CD11b, GR1, Ly6C, CD11c, MHCII, CD64, PDCA-1, F4/80, CD163, CD68, CD86, CD206. DAPI was used to exclude dead cells. Multiparameter analysis was performed on a LSR Fortessa (BD) and bi-dimensional dot plots were generated using FlowJo software.

### Lipid mediator (LM) metabololipidomics

Colons containing stool were obtained after DSS-colitis at day 5 and day 12 and were snap frozen in liquid nitrogen and stored at −80°C for liquid chromatography-tandem mass spectrometry (LC-MS/MS) metabololipidomics analyses. Prior to sample extraction, ice-cold methanol including d_8_-5-HETE, d_5_-RvD2, d_5_-LXA_4_, d_4_-LTB_4_, and d_4_-PGE_2_ (500 pg each; Cayman Chemical) were added to facilitate quantification. Next, colons were homogenized manually with a tissue grinder manifold kit on ice for 1 minute. After, all samples were kept at −80°C for 45 min to allow protein precipitation and then spun at 1000g at 4°C for 5 min. After centrifugation, each supernatant was rapid acidified (<30 s) to ∼pH 3.5 and then loaded onto solid-phase extraction (SPE) Isolute C18 SPE 3-mL, 100 mg cartridges (Biotage). Before elution, LM bound to the SPE matrix were neutralized with ddH_2_O and washed with 3ml of hexane. LMs were eluted with 3ml of methyl formate. Methyl formate fractions from the SPE were brought to dryness under a stream of nitrogen and resuspended with 50μL in 1:1 methanol:water before injection into a liquid chromatography-tandem mass spectrometry system consisting of a QTrap 5500 (AB Sciex) equipped with a Shimadzu LC-20AD HPLC (Tokyo, Japan). A Poroshell 120 EC-18 column (100 mm × 4.6 mm × 2.7 μm; Agilent Technologies) was kept in a column oven maintained at 50°C, and LMs were eluted in a gradient of methanol/water/acetic acid from 55:45:0.01 (v/v/v) to 100:0:0.01 at a flow rate of 0.5 ml/min. To monitor and quantify the amounts of lipid mediators of interest, multiple reaction monitoring (MRM) was used with MS/MS matching signature ion fragments for each molecule with at least six diagnostic ions; ∼0.1 pg limits of detection. Principal Component Analysis (PCA) was performed with SIMCA software, version 10.0.3, MKS Data Analytics Solutions, Sweden.

#### *In vivo* SPMs treatment

Specialized pro-resolving mediator treatment was performed using a mix of SPMs purchased from Cayman Chemical Ann Arbor, Michigan (15(*R*)-LXA_4_ 90415; RvD1 10012554; RvD2 10007279; RvD4 13835; RvE2 (13827)) a mixture of 100ng of each SPM was prepared to a final total amount of 500ng of SPMs. The mixture was treated with a gentle stream of nitrogen to evaporate the methanol carrier and resuspended in ethanol. Treatment consisted of 500ng of SPM-mixture in 100μL saline per injection per animal i.p. Vehicle treatment was 0.1% of ethanol in 100 μL of saline. Mice were SPM or vehicle dosed every other day, from day 5 to day 12.

## Acknowledgments

We thank the members of the Yaffe lab for helpful advice and discussions. We acknowledge the MIT BioMicro Center and Genomics Facility, the Koch Institute’s Robert A. Swanson (1969) Biotechnology Center for technical support, specifically the Preclinical Modeling Facility, the Flow Cytometry Core Facility, and the Hope Babette Tang (1983) Histology Facility. NIH BioArt was used to create the graphical abstract.

## Author Contributions

LS-L, CNS and MBY conceived the project, LS-L, XY, XR, TE, YWK, CNS and MBY designed the research studies, LS-L, XY, XR, and YWK conducted experiments, HAB, LS-L, XY, XR, TE, PF, YWK, EJV, CCK, CNS and MBY analyzed data, HAB, LS-L, XY, PF and MBY wrote the manuscript and all authors performed manuscript revisions.

## Use of Artificial Intelligence

The authors used ChatGPTv5.5 and Claude (Anthropic) between June and October 2026 to assist with revising portions of the manuscript text, internal consistency checking, and verification of references and reported values against primary sources. All study concepts, analyses, and interpretations are the authors’ own; the authors reviewed and revised all content and take full responsibility for the integrity and accuracy of the manuscript.

## Data Sharing Statement

A data file for all graphs in this manuscript is provided in the supplemental documents.

