## Supplemental Tables for "MAPKAP Kinase-2 signaling enhances the resolution of colonic inflammation through specialized pro-resolving mediators"

### Supplemental Table 1. Colonic cytokine and chemokine levels following DSS-induced colitis: summary statistics.

Values are mean ± SEM (pg/mL) from a multiplex cytokine/chemokine panel measured in colon tissue at the indicated day of DSS colitis; n = animals per group. Control is WT for the global knockout and Fl/Fl for the myeloid-restricted knockout.

| Analyte | Day | Control (WT or Fl/Fl) | MK2 KO (MK2KO or LysM-KO) |
|---|---|---|---|
| ***Global MK2 knockout (whole-body): WT vs MK2KO*** | | | |
| **IL-13** | 0 | 0.00 ± 0.00 (n=3) | 0.00 ± 0.00 (n=3) |
| | 5 | 8.39 ± 0.58 (n=3) | 3.52 ± 2.12 (n=6) |
| | 9 | 8.21 ± 0.28 (n=4) | 3.42 ± 1.73 (n=5) |
| **IL-10** | 0 | 10.95 ± 1.51 (n=3) | 21.04 ± 7.78 (n=3) |
| | 5 | 96.94 ± 9.26 (n=3) | 43.51 ± 21.84 (n=7) |
| | 9 | 34.81 ± 6.73 (n=4) | 10.80 ± 6.22 (n=5) |
| **M-CSF** | 0 | 1.96 ± 1.00 (n=3) | 0.98 ± 0.50 (n=3) |
| | 5 | 6.48 ± 0.81 (n=3) | 2.52 ± 0.96 (n=6) |
| | 9 | 5.92 ± 0.61 (n=4) | 4.09 ± 0.71 (n=5) |
| **IL-6** | 0 | 0.60 ± 0.10 (n=3) | 0.43 ± 0.22 (n=3) |
| | 5 | 49.05 ± 29.09 (n=3) | 239.31 ± 105.46 (n=6) |
| | 9 | 24.59 ± 10.36 (n=4) | 636.79 ± 338.88 (n=5) |
| **Eotaxin** | 0 | 43.35 ± 4.27 (n=3) | 71.90 ± 17.61 (n=3) |
| | 5 | 121.67 ± 17.43 (n=3) | 193.75 ± 44.43 (n=6) |
| | 9 | 72.61 ± 7.22 (n=4) | 308.44 ± 86.91 (n=5) |
| **G-CSF** | 0 | 0.91 ± 0.18 (n=3) | 0.79 ± 0.33 (n=3) |
| | 5 | 26.73 ± 9.97 (n=3) | 81.72 ± 44.90 (n=6) |
| | 9 | 44.06 ± 17.76 (n=4) | 538.87 ± 245.37 (n=5) |
| **IL-17** | 0 | 0.18 ± 0.18 (n=3) | 1.73 ± 0.60 (n=3) |
| | 5 | 1.56 ± 0.25 (n=3) | 2.12 ± 0.37 (n=6) |
| | 9 | 2.37 ± 0.41 (n=4) | 14.57 ± 7.66 (n=5) |
| **LIF** | 0 | 1.30 ± 0.20 (n=3) | 1.28 ± 0.27 (n=3) |
| | 5 | 3.63 ± 0.56 (n=3) | 6.75 ± 1.81 (n=6) |
| | 9 | 2.79 ± 0.49 (n=4) | 14.76 ± 6.74 (n=5) |
| ***Myeloid-restricted MK2 knockout (LysM-Cre): Fl/Fl vs LysM-KO*** | | | |
| **IL-6** | 5 | 62.81 ± 29.41 (n=3) | 101.07 ± 51.96 (n=3) |
| | 9 | 39.48 ± 27.68 (n=4) | 208.13 ± 88.37 (n=3) |
| **Eotaxin** | 5 | 156.60 ± 16.70 (n=3) | 151.73 ± 3.14 (n=3) |
| | 9 | 156.72 ± 26.32 (n=4) | 224.08 ± 15.62 (n=3) |
| **IL-17** | 5 | 5.95 ± 0.92 (n=3) | 5.52 ± 1.19 (n=3) |
| | 9 | 6.36 ± 0.60 (n=4) | 10.37 ± 1.85 (n=3) |
| **KC** | 5 | 66.50 ± 3.90 (n=3) | 121.13 ± 29.66 (n=3) |
| | 9 | 44.25 ± 14.23 (n=4) | 135.79 ± 39.80 (n=3) |
| **G-CSF** | 5 | 41.46 ± 5.15 (n=3) | 81.03 ± 24.77 (n=3) |
| | 9 | 55.14 ± 36.19 (n=4) | 316.56 ± 106.99 (n=3) |

### Supplemental Table 2. Colonic cytokine and chemokine levels following DSS-induced colitis: individual replicate values.

Individual replicate values (pg/mL) underlying Table 1. Control is WT for the global knockout and Fl/Fl for the myeloid-restricted knockout.

| Analyte | Day | Control (WT or Fl/Fl) | MK2 KO (MK2KO or LysM-KO) |
|---|---|---|---|
| *Global MK2 knockout (whole-body): WT vs MK2KO* | | | |
| **IL-13** | 0 | 0, 0, 0 | 0, 0, 0 |
| | 5 | 9.26, 8.63, 7.29 | 8.63, 11.58, 0, 0, 0, 0.92 |
| | 9 | 7.86, 8.18, 9.01, 7.8 | 9.52, 5.04, 0.76, 1.02, 0.78 |
| **IL-10** | 0 | 9.69, 9.2, 13.95 | 9.42, 35.8, 17.9 |
| | 5 | 78.93, 109.69, 102.19 | 86.15, 141.48, 76.43, 0, 0, 0, 0.53 |
| | 9 | 24.06, 53.15, 36.63, 25.42 | 29.2, 22.37, 0, 0.64, 1.78 |
| **M-CSF** | 0 | 3.25, 0, 2.62 | 1.29, 1.64, 0 |
| | 5 | 4.92, 6.86, 7.65 | 5.3, 5.49, 0, 1.35, 0.64, 2.36 |
| | 9 | 5.3, 7.18, 6.69, 4.53 | 5.67, 5.67, 3.94, 3.12, 2.06 |
| **IL-6** | 0 | 0.8, 0.52, 0.47 | 0.09, 0.85, 0.36 |
| | 5 | 10.39, 106.02, 30.73 | 75.02, 597.64, 13.65, 104.7, 104.05, 540.82 |
| | 9 | 41.7, 9.28, 4.2, 43.19 | 6.19, 29.52, 1624.75, 1273.12, 250.38 |
| **Eotaxin** | 0 | 51.3, 42.07, 36.69 | 41.88, 102.85, 70.96 |
| | 5 | 92.65, 152.92, 119.45 | 117.3, 350.2, 276.5, 175.99, 46.3, 196.19 |
| | 9 | 51.83, 80.47, 83.98, 74.15 | 154.83, 87.61, 474.44, 533.95, 291.38 |
| **G-CSF** | 0 | 0.64, 0.84, 1.26 | 0.79, 1.36, 0.23 |
| | 5 | 11.24, 45.34, 23.62 | 29.47, 294.55, 4.03, 20.72, 35.81, 105.77 |
| | 9 | 16.54, 90.44, 53.31, 15.95 | 6.08, 196.39, 1055.22, 1204.1, 232.58 |
| **IL-17** | 0 | 0, 0.55, 0 | 0.95, 2.92, 1.33 |
| | 5 | 2.06, 1.25, 1.36 | 3.5, 2.71, 2.18, 1.71, 0.83, 1.8 |
| | 9 | 1.57, 3.04, 3.12, 1.75 | 2.13, 4.67, 21.25, 41.82, 3 |
| **LIF** | 0 | 1.55, 0.91, 1.45 | 1.09, 1.81, 0.93 |
| | 5 | 2.74, 4.66, 3.48 | 3.73, 12.83, 2.98, 4.22, 4.73, 11.98 |
| | 9 | 1.76, 4.07, 2.98, 2.37 | 2.52, 3.23, 33.87, 28.26, 5.91 |
| *Myeloid-restricted MK2 knockout (LysM-Cre): Fl/Fl vs LysM-KO* | | | |
| **IL-6** | 5 | 16.79, 117.55, 54.09 | 48.65, 49.57, 205 |
| | 9 | 3.02, 3.44, 31.37, 120.09 | 191.17, 368.96, 64.25 |
| **Eotaxin** | 5 | 143.18, 189.79, 136.82 | 157.98, 148.07, 149.13 |
| | 9 | 102.57, 132.81, 166.06, 225.42 | 243.54, 235.52, 193.19 |
| **IL-17** | 5 | 4.3, 6.05, 7.49 | 4.7, 3.98, 7.87 |
| | 9 | 6.44, 5.03, 6.04, 7.92 | 13.58, 10.38, 7.16 |
| **KC** | 5 | 65.47, 60.32, 73.71 | 76.77, 177.42, 109.2 |
| | 9 | 36.95, 25.56, 28.19, 86.3 | 132.8, 206.16, 68.4 |
| **G-CSF** | 5 | 40.82, 50.69, 32.88 | 103.37, 31.57, 108.15 |
| | 9 | 16.63, 1.29, 41.89, 160.76 | 362.73, 474.43, 112.53 |

**Supplemental Table 3. Abbreviations and complete stereochemical assignments of specialized pro-resolving mediators (SPMs) monitored in this study**

| Abbreviation | Full name | Complete stereochemical name | Detected in this dataset |
|---|---|---|---|
| **D-series resolvins (from docosahexaenoic acid)** | | | |
| **RvD1** | Resolvin D1 | 7S,8R,17S-trihydroxydocosa-4Z,9E,11E,13Z,15E,19Z-hexaenoic acid | Yes |
| **RvD2** | Resolvin D2 | 7S,16R,17S-trihydroxydocosa-4Z,8E,10Z,12E,14E,19Z-hexaenoic acid | Yes |
| **RvD3** | Resolvin D3 | 4S,11R,17S-trihydroxydocosa-5Z,7E,9E,13Z,15E,19Z-hexaenoic acid | No |
| **RvD4** | Resolvin D4 | 4S,5R,17S-trihydroxydocosa-6E,8E,10Z,13Z,15E,19Z-hexaenoic acid | Yes |
| **RvD5** | Resolvin D5 | 7S,17S-dihydroxydocosa-4Z,8E,10Z,13Z,15E,19Z-hexaenoic acid | Yes |
| **RvD6** | Resolvin D6 | 4S,17S-dihydroxydocosa-5E,7Z,10Z,13Z,15E,19Z-hexaenoic acid | No |
| **AT-RvD1** | Aspirin-triggered Resolvin D1 | 7S,8R,17R-trihydroxydocosa-4Z,9E,11E,13Z,15E,19Z-hexaenoic acid | Yes |
| **AT-RvD3** | Aspirin-triggered Resolvin D3 | 4S,11R,17R-trihydroxydocosa-5Z,7E,9E,13Z,15E,19Z-hexaenoic acid | Yes |
| **Protectins (from docosahexaenoic acid)** | | | |
| **PD1 / NPD1** | Protectin D1 / Neuroprotectin D1 | 10R,17S-dihydroxydocosa-4Z,7Z,11E,13E,15Z,19Z-hexaenoic acid | Yes |
| **AT-PD1** | Aspirin-triggered Protectin D1 | 10R,17R-dihydroxydocosa-4Z,7Z,11E,13E,15Z,19Z-hexaenoic acid | Yes |
| **10S,17S-diHDHA** | 10S,17S-dihydroxydocosahexaenoic acid | 10S,17S-dihydroxydocosa-4Z,7Z,11E,13Z,15E,19Z-hexaenoic acid | Yes |
| **22-OH-PD1** | ω-22-hydroxy Protectin D1 | 10R,17S,22-trihydroxydocosa-4Z,7Z,11E,13E,15Z,19Z-hexaenoic acid | Yes |
| **Maresins (from docosahexaenoic acid)** | | | |
| **Maresin 1 (MaR1)** | Maresin 1 | 7R,14S-dihydroxydocosa-4Z,8E,10E,12Z,16Z,19Z-hexaenoic acid | No |
| **7S,14S-diHDHA** | 7-epi-Maresin 1 | 7S,14S-dihydroxydocosa-4Z,8E,10Z,12E,16Z,19Z-hexaenoic acid | No |
| **4S,14S-diHDHA** | 4S,14S-dihydroxydocosahexaenoic acid | 4S,14S-dihydroxydocosahexaenoic acid | No |
| **E-series resolvins (from eicosapentaenoic acid)** | | | |
| **RvE1** | Resolvin E1 | 5S,12R,18R-trihydroxyeicosa-6Z,8E,10E,14Z,16E-pentaenoic acid | Yes |
| **RvE2** | Resolvin E2 | 5S,18R-dihydroxyeicosa-6E,8Z,11Z,14Z,16E-pentaenoic acid | Yes |
| **RvE3** | Resolvin E3 | 17R,18R/17S,18S-dihydroxyeicosa-5Z,8Z,11Z,13E,15E-pentaenoic acid | Yes |
| **Lipoxins (from arachidonic acid)** | | | |
| **LXA4** | Lipoxin A4 | 5S,6R,15S-trihydroxyeicosa-7E,9E,11Z,13E-tetraenoic acid | Yes |
| **LXB4** | Lipoxin B4 | 5S,14R,15S-trihydroxyeicosa-6E,8Z,10E,12E-tetraenoic acid | Yes |
| **5S,15S-diHETE** | 5S,15S-dihydroxyeicosatetraenoic acid | 5S,15S-dihydroxyeicosa-6E,8Z,11Z,13E-tetraenoic acid | Yes |
| **AT-LXA4** | Aspirin-triggered Lipoxin A4 (15-epi-LXA4) | 5S,6R,15R-trihydroxyeicosa-7E,9E,11Z,13E-tetraenoic acid | Yes |
| **AT-LXB4** | Aspirin-triggered Lipoxin B4 (15-epi-LXB4) | 5S,14R,15R-trihydroxyeicosa-6E,8Z,10E,12E-tetraenoic acid | Yes |

### Supplemental Table 4. Targeted lipid mediator metabololipidomics of murine colon tissue after DSS-induced colitis: specialized pro-resolving mediators

Values are concentrations (pg/500 mg tissue) measured by targeted LC-MS/MS (Qtrap 5500) in WT and MK2-deficient (KO) mouse colons at days 5 and 12 of DSS-induced colitis (n=4 WT animals, n=3 KO animals per timepoint). 'ND' = not detected (below the assay's limit of detection, ~0.1 pg).

| Analyte | Q1 | Q3 | WT D5 | WT D12 | KO D5 | KO D12 |
|---|---|---|---|---|---|---|
| **D-series resolvins** | | | | | | |
| **RvD1** | 375 | 121 | 8.1 ± 1.2 (n=4) | 182.8 ± 91.1 (n=4) | 0.9 ± 0.0 (n=3) | 1.9 ± 0.0 (n=3) |
| **RvD2** | 375 | 215 | 266.0 ± 59.3 (n=4) | 990.8 ± 259.9 (n=4) | 273.7 ± 60.1 (n=3) | 201.2 ± 72.9 (n=3) |
| **RvD3** | 375 | 181 | ND | ND | ND | ND |
| **RvD4** | 375 | 101 | 11.1 ± 5.5 (n=4) | 84.2 ± 15.2 (n=4) | 4.2 ± 1.6 (n=3) | 18.4 ± 2.1 (n=3) |
| **RvD5** | 359 | 199 | 69.6 ± 14.4 (n=4) | 121.5 ± 14.7 (n=4) | 77.3 ± 52.1 (n=3) | 12.6 ± 4.9 (n=3) |
| **RvD6** | 359 | 101 | ND | ND | ND | ND |
| **AT-RvD1** | 375 | 121 | 50.4 ± 24.7 (n=4) | 76.1 ± 19.3 (n=4) | 11.7 ± 4.1 (n=3) | 19.9 ± 3.0 (n=3) |
| **AT-RvD3** | 375 | 181 | 234.6 ± 95.9 (n=4) | 120.8 ± 26.2 (n=4) | 8.6 ± 2.4 (n=3) | 17.0 ± 4.7 (n=3) |
| **Protectins** | | | | | | |
| **PD1** | 359 | 153 | 9.8 ± 2.0 (n=4) | 7.7 ± 2.6 (n=4) | 4.3 ± 2.2 (n=3) | 2.9 ± 0.5 (n=3) |
| **AT-PD1** | 359 | 153 | 7.6 ± 2.6 (n=4) | 23.7 ± 7.8 (n=4) | 8.7 ± 1.9 (n=3) | 12.2 ± 2.8 (n=3) |
| **10S,17S-diHDHA** | 359 | 153 | 85.4 ± 31.0 (n=4) | 52.5 ± 6.1 (n=4) | 31.3 ± 6.0 (n=3) | 33.6 ± 2.6 (n=3) |
| **22-OH-PD1** | 375 | 153 | 904.6 ± 285.5 (n=4) | 3055.4 ± 786.6 (n=4) | 491.2 ± 179.2 (n=3) | 1102.0 ± 124.8 (n=3) |
| **Maresins** | | | | | | |
| **Maresin 1** | 359 | 221 | ND | ND | ND | ND |
| **7S,14S-diHDHA** | 359 | 221 | ND | ND | ND | ND |
| **4S,14S-diHDHA** | 359 | 101 | ND | ND | ND | ND |
| **E-series resolvins** | | | | | | |
| **RvE1** | 349 | 195 | 44.2 ± 9.8 (n=4) | 29.2 ± 7.3 (n=4) | 9.2 ± 2.0 (n=3) | 25.7 ± 7.9 (n=3) |
| **RvE2** | 333 | 199 | 41.7 ± 27.2 (n=4) | 170.8 ± 32.8 (n=4) | 8.9 ± 3.2 (n=3) | 17.0 ± 5.8 (n=3) |
| **RvE3** | 333 | 201 | 74.6 ± 26.2 (n=4) | 117.3 ± 28.1 (n=4) | 29.7 ± 4.4 (n=3) | 25.3 ± 10.6 (n=3) |
| **Lipoxins** | | | | | | |
| **LXA4** | 351 | 115 | 411.9 ± 291.2 (n=4) | 1547.8 ± 728.2 (n=4) | 44.1 ± 23.1 (n=3) | 94.7 ± 46.7 (n=3) |
| **LXB4** | 351 | 221 | 788.9 ± 277.7 (n=4) | 4531.4 ± 2070.5 (n=4) | 141.3 ± 11.9 (n=3) | 683.4 ± 182.5 (n=3) |
| **5S,15S-diHETE** | 335 | 115 | 590.8 ± 328.3 (n=4) | 777.9 ± 120.6 (n=4) | 406.5 ± 212.0 (n=3) | 416.7 ± 132.3 (n=3) |
| **AT-LXA4** | 351 | 115 | 182.3 ± 45.1 (n=4) | 859.1 ± 190.3 (n=4) | 89.4 ± 25.0 (n=3) | 234.3 ± 56.7 (n=3) |
| **AT-LXB4** | 351 | 221 | 143.7 ± 94.5 (n=4) | 276.3 ± 33.1 (n=4) | 34.0 ± 2.8 (n=3) | 99.8 ± 25.3 (n=3) |

### Supplemental Table 5. Targeted lipid mediator metabololipidomics of murine colon tissue after DSS-induced colitis: pro-inflammatory eicosanoids and pathway markers/precursor fatty acids

Values are concentrations (pg/500 mg tissue) measured by targeted LC-MS/MS (Qtrap 5500) in WT and MK2-deficient (KO) mouse colons at day 5 and day 12 of DSS-induced colitis (n=4 WT animals, n=3 KO animals per timepoint). 'ND' = not detected (below the assay's limit of detection, ~0.1 pg).

| Analyte | Q1 | Q3 | WT D5 | WT D12 | KO D5 | KO D12 |
|---|---|---|---|---|---|---|
| **Pro-inflammatory eicosanoids** | | | | | | |
| **LTB4** | 335 | 195 | 284.8 ± 129.7 (n=4) | 407.7 ± 145.3 (n=4) | 427.8 ± 81.1 (n=3) | 63.9 ± 16.6 (n=3) |
| **20-COOH-LTB4** | 365 | 195 | ND | ND | ND | ND |
| **5S,12S-diHETE** | 335 | 195 | 90.3 ± 18.9 (n=4) | 206.1 ± 76.1 (n=4) | 130.9 ± 10.3 (n=3) | 59.8 ± 4.5 (n=3) |
| **PGD2** | 351 | 233 | 1702.7 ± 241.6 (n=4) | 3200.1 ± 534.4 (n=4) | 823.9 ± 158.4 (n=3) | 3879.8 ± 1005.9 (n=3) |
| **PGE2** | 351 | 189 | 1815.7 ± 239.7 (n=4) | 3529.0 ± 430.2 (n=4) | 1086.6 ± 12.5 (n=3) | 2363.5 ± 400.8 (n=3) |
| **PGF2a** | 353 | 193 | 1761.1 ± 268.8 (n=4) | 3375.0 ± 540.4 (n=4) | 562.5 ± 147.1 (n=3) | 3524.3 ± 1031.5 (n=3) |
| **TXB2** | 369 | 169 | 703.2 ± 167.2 (n=4) | 1269.5 ± 253.1 (n=4) | 312.5 ± 57.8 (n=3) | 1004.5 ± 125.8 (n=3) |
| **Pathway markers / precursor fatty acids** | | | | | | |
| **17-HDHA** | 343 | 245 | 3923.7 ± 1418.7 (n=4) | 857.7 ± 236.5 (n=4) | 923.6 ± 45.5 (n=3) | 1058.1 ± 145.8 (n=3) |
| **14-HDHA** | 343 | 205 | 3016.3 ± 982.8 (n=4) | 500.9 ± 28.6 (n=4) | 756.1 ± 72.7 (n=3) | 754.6 ± 81.4 (n=3) |
| **7-HDHA** | 343 | 141 | 523.3 ± 171.2 (n=4) | 278.2 ± 21.6 (n=4) | 212.8 ± 21.0 (n=3) | 277.9 ± 26.9 (n=3) |
| **4-HDHA** | 343 | 101 | 590.9 ± 305.9 (n=4) | 305.7 ± 26.4 (n=4) | 189.0 ± 10.9 (n=3) | 371.4 ± 36.9 (n=3) |
| **DHA** | 327 | 283 | 101442.9 ± 33419.2 (n=4) | 70436.1 ± 8059.1 (n=4) | 24562.5 ± 3358.9 (n=3) | 44609.6 ± 1457.1 (n=3) |
| **18-HEPE** | 317 | 259 | 1454.1 ± 263.2 (n=4) | 1463.0 ± 186.1 (n=4) | 407.2 ± 51.4 (n=3) | 1185.2 ± 162.6 (n=3) |
| **15-HEPE** | 317 | 219 | 2303.1 ± 577.8 (n=4) | 1733.1 ± 155.4 (n=4) | 754.0 ± 110.9 (n=3) | 1283.7 ± 244.2 (n=3) |
| **12-HEPE** | 317 | 179 | 2430.6 ± 903.3 (n=4) | 430.6 ± 26.9 (n=4) | 709.6 ± 68.5 (n=3) | 457.5 ± 70.9 (n=3) |
| **5-HEPE** | 317 | 115 | 1058.1 ± 385.1 (n=4) | 592.8 ± 131.1 (n=4) | 333.3 ± 38.7 (n=3) | 610.1 ± 24.5 (n=3) |
| **EPA** | 301 | 257 | 212520.2 ± 68079.7 (n=4) | 154266.2 ± 18855.9 (n=4) | 54359.4 ± 9607.3 (n=3) | 82210.3 ± 5499.6 (n=3) |
| **15-HETE** | 319 | 219 | 1262.2 ± 441.8 (n=4) | 294.3 ± 32.1 (n=4) | 169.8 ± 12.5 (n=3) | 634.8 ± 106.9 (n=3) |
| **12-HETE** | 319 | 179 | 1442.3 ± 536.0 (n=4) | 246.9 ± 22.7 (n=4) | 421.1 ± 40.7 (n=3) | 271.5 ± 42.1 (n=3) |
| **5-HETE** | 319 | 115 | 1010.7 ± 311.9 (n=4) | 761.2 ± 141.7 (n=4) | 511.1 ± 10.6 (n=3) | 645.9 ± 28.2 (n=3) |
| **AA** | 303 | 257 | 196060.0 ± 83047.7 (n=4) | 145311.3 ± 21367.8 (n=4) | 47738.2 ± 8629.0 (n=3) | 67202.9 ± 4308.0 (n=3) |
